# Atg8 orchestrates stress-responsive chromatin programs across immunity and metabolism

**DOI:** 10.64898/2026.05.22.727304

**Authors:** Kevin P. Kelly, Navyashree A. Ramesh, Sunidhi Ranganathan, Aditi Madan, Shannon Marschall, Robert L. Unckless, Akhila Rajan

**Author notes:** Correspondence (K.P.K.), (A.R.).

## Abstract

Organisms must coordinate transcriptional responses to immune and metabolic stress, often within the same tissue. In Drosophila and mammals, adipose tissue integrates these signals by mounting antimicrobial defense during acute infection and remodeling lipid metabolism under chronic nutrient surplus. How one cell-biological system supports both functions, and through what molecular machinery, remains incompletely understood. Atg8/LC3, classically defined by canonical autophagy, has emerging non-canonical roles in nuclear gene regulation, raising the possibility that it contributes to stress-coordinated transcription beyond cargo turnover. Using unbiased CUT&RUN in adult Drosophila nuclei, we find that endogenous Atg8 exhibits broad chromatin occupancy at immune, metabolic, and autophagy loci, and accumulates in nuclei under prolonged high-sugar diet (HSD) and acute Gram-positive infection. We identify two conserved Atg8-interacting motifs (AIMs) within the Rel homology domain of NF-κB/Dif. Flies carrying CRISPR-engineered AIM-mutant Dif are highly susceptible to both infection and chronic HSD, establishing a physiological requirement for intact Dif AIMs. AIM-mutant Dif shows impaired infection-induced nuclear accumulation, suggesting that Atg8 contributes to both Dif cytoplasmic-to-nuclear shuttling and nuclear function. Unbiased comparison of Atg8 chromatin occupancy across HSD and infection further reveals shared and divergent motif grammar, positioning Atg8 as a stress-responsive chromatin cofactor for immune and metabolic transcription. Together, these findings expand the functional landscape of Atg8/LC3 beyond canonical autophagy and reveal that autophagy machinery contributes to stress-specific transcriptional complex assembly. AIM/LIR-mediated interactions, exemplified by Dif, represent one such interface, while additional mechanisms likely underlie Atg8’s broader chromatin engagement at loci enriched for transcription factor motifs whose cognate factors lack known AIM/LIRs. We propose that Atg8/LC3-mediated coordination of immune and metabolic transcription is a general principle by which cells integrate diverse stress signals, with implications for obesity, chronic inflammation, and other disease states in which immune and metabolic dysregulation converge.

**GRAPHICAL ABSTRACT:** 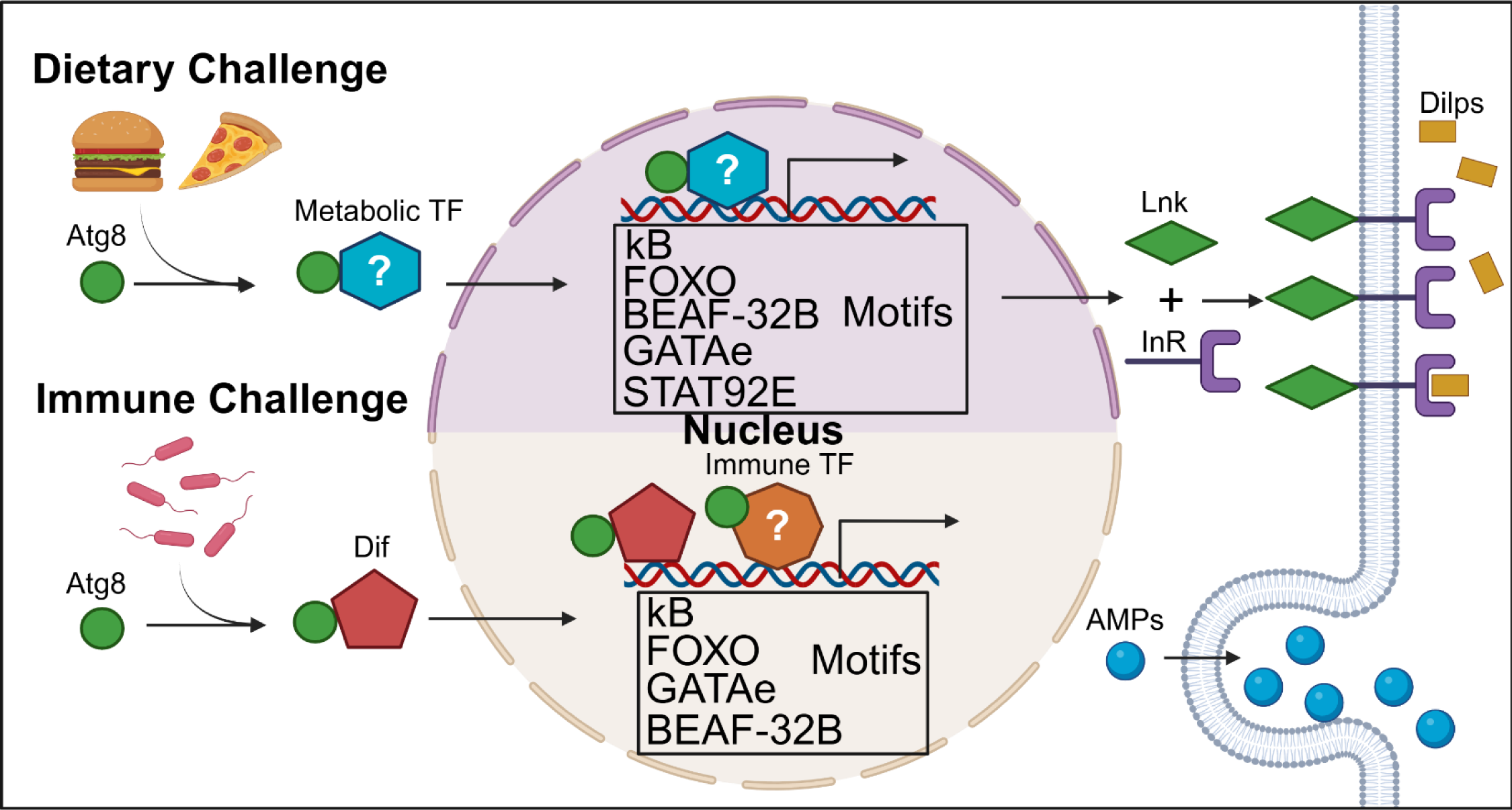

- Stress drives Atg8 into nuclei, where it occupies immune and metabolic chromatin.
- Two conserved AIMs in NF-κB/Dif bind Atg8 and enable Dif nuclear entry.
- AIM-mutant Dif flies are highly susceptible to infection and chronic high-sugar diet.
- Atg8 occupies stress-related motifs on prolonged HSD and acute infection.

## INTRODUCTION

Small ubiquitin-like modifiers profoundly shape eukaryotic biology. Ubiquitin, the founding member of this structurally conserved family, was originally characterized for its role in directing substrate proteins to proteasomal degradation[1]. Subsequent work has reshaped this view considerably. Ubiquitin is now recognized as a regulator of transcription in its own right — through monoubiquitination of histone H2A and H2B at chromatin[2–4], through ubiquitin-mediated control of NF-κB activation[5], and through assembly of polyubiquitin chains that orchestrate signaling complexes well beyond proteolytic destinations[6, 7]. This expanded view of ubiquitin biology has revealed that a single conserved modifier can carry diverse, context-dependent regulatory information, and has motivated parallel inquiry into the broader family of ubiquitin-like proteins that operate alongside it.

Atg8 (LC3/GABARAP in mammals), a conserved ubiquitin-like protein, exemplifies this conceptual expansion. Atg8 is best known for its canonical role in macroautophagy, where covalent conjugation to phosphatidylethanolamine (PE) anchors Atg8 onto nascent autophagosomal membranes and supports cargo selection, autophagosome maturation, and lysosomal turnover during nutrient scarcity[8–11]. LC3 phosphorylation further regulates the directional trafficking of autophagosomes toward perinuclear lysosomal compartments to ensure efficient cargo turnover[12]. Over the past decade, however, a substantial body of work has revealed an expanding repertoire of non-canonical functions for the Atg8/LC3 family that extend well beyond classical macroautophagic degradation[13–16]. The Atg8 conjugation machinery operates at single-membrane organelles in LC3-associated phagocytosis (LAP) and LC3-associated endocytosis (LANDO), where it mediates cargo turnover and receptor recycling, including microglial clearance of amyloid-β[13, 17, 18]. Atg8/LC3 directs RNA-binding proteins and microRNAs into secreted extracellular vesicles through LC3-dependent EV loading and secretion (LDELS)[19, 20], supports the unconventional secretion of leaderless cytokines such as IL-1β through secretory autophagosomes[21, 22], and is hijacked by enveloped viruses for membrane acquisition during egress[23]. The Atg8/LC3 system also modulates innate immune signaling, regulating pattern recognition receptors and interferon production during infection[24]. In our own recent work, Atg8 mediates the unconventional secretion of the fly adipokine Upd2 and of human leptin from adipocytes through the LDELS pathway[25]. Collectively, these findings establish Atg8/LC3 as a versatile component of cellular trafficking and signaling networks whose biological deployment extends well beyond cargo degradation.

Within this expanding landscape, the nuclear and transcriptional roles of Atg8 remain comparatively less explored. Atg8 interacts with the transcription factor Sequoia to regulate autophagy gene expression in *Drosophila*[*26*], undergoes deacetylation-driven nucleocytoplasmic shuttling under starvation[27], and engages the transcription factors LMX1A/B to regulate autophagy gene transcription during mammalian neuronal development[28]. The non-canonical functions of Atg8 are unified by a single biochemical logic: Atg8 lacks intrinsic DNA-binding activity and engages partner proteins instead through the short linear Atg8-interacting motif (AIM in flies, LIR in mammals; consensus W/F/Y-x-x-L/V/I), a degenerate and broadly distributed sequence that likely permits recruitment of a substantially larger set of partner proteins than has so far been catalogued[29, 30]. Yet the reported nuclear functions of Atg8 have been largely circumscribed to the regulation of autophagy genes themselves, leaving open the question of whether Atg8 engages chromatin more broadly — and whether such engagement could coordinate transcriptional outputs across distinct physiological contexts.

Adipose tissue provides an experimentally tractable setting in which to address this question. The adult *Drosophila* fat body — like mammalian adipose tissue — is a stress-responsive endocrine organ that integrates information from metabolic, immune, and neuroendocrine signals to produce coordinated systemic outputs[31–33]. Two stressors place opposing demands on this integrative function. Chronic high-sugar diet (HSD), a well-established obesogenic challenge in flies and mammals, disrupts fat body lipid homeostasis, alters phospholipid composition (including PE, the very lipid to which Atg8 is conjugated), and reshapes the metabolic transcriptome over the course of weeks[34–38]. Gram-positive bacterial infection, by contrast, demands an acute reorganization of adipocyte output toward humoral defense, including the rapid induction of antimicrobial peptides through Toll receptor signaling and the NF-κB transcription factor Dif[39–44]. Importantly, these two challenges are not biologically independent: recent work has shown that chronic high-sugar diet itself increases susceptibility to bacterial infection in *Drosophila*[*45*], and chronic infection imposes substantial metabolic costs on the organism[46], reinforcing the case for examining metabolic and immune stress responses through a shared cellular machinery in the same tissue. Consistent with this view, bidirectional crosstalk between NF-κB-family transcription factors and the autophagy machinery has been documented in other contexts, including transcriptional control of *Atg1* expression during developmental autophagy[47]. Because Atg8 nuclear accumulation has been reported under both nutrient deprivation and immune challenge, and because the AIM motif is broadly distributed across the transcription factor landscape, we reasoned that Atg8 may participate in transcriptional regulation in adipose tissue across multiple stress contexts — and that the regulatory output it supports may be shaped by the partner factors available within each particular physiological state.

Here, we test this possibility directly. Using unbiased CUT&RUN profiling of endogenous Atg8 in adult *Drosophila* nuclei; an inducible degron strategy for adult-stage Atg8 depletion that bypasses developmental lethality; CRISPR-engineered AIM-mutant alleles of the NF-κB transcription factor Dif; and matched comparisons across acute gram-positive infection and chronic HSD, we ask three connected questions. First, where does Atg8 engage chromatin under baseline conditions, and at what motifs is its occupancy enriched? Second, how is Atg8 chromatin engagement reshaped by infection versus chronic high-sugar diet? And third, through what molecular interactions does Atg8 contribute to transcriptional output at the loci it engages? We find that Atg8 occupies promoter-proximal chromatin broadly across the genome, with enrichment at autophagy, immune, and metabolic loci; that its nuclear accumulation and chromatin redeployment are coordinately reorganized by stress, with substantial context-specific redistribution between immune and metabolic targets; and that Atg8–Dif interaction via two conserved AIMs within Dif’s Rel Homology domain is required for productive Dif chromatin engagement, antimicrobial peptide induction, and organismal resilience under both infection and chronic HSD. Together, these findings position Atg8 not as an autophagy effector that occasionally enters the nucleus, but as a stress-responsive chromatin-associated cofactor whose context-sensitive deployment integrates immune and metabolic stress responses in adipose tissue.

## RESULTS

### Characterizing Atg8 chromatin binding at transcription sites in the basal state

Recent studies have identified Atg8/LC3 as a regulator of gene expression in both *Drosophila melanogaster* and humans, particularly in the regulation of autophagy-related genes[26, 28]. Given the broad nature of the Atg8-interacting motif (AIM; referred to as the LC3-interacting region, or LIR, in humans), we sought to identify additional transcriptional regulators that may associate with Atg8. To investigate the role of Atg8 in transcriptional regulation, we performed Cleavage Under Targets and Release Using Nuclease (CUT&RUN) using anti-Atg8a antibodies to identify chromatin regions occupied by Atg8. Because Atg8 lacks a canonical DNA-binding domain, we employed a modified low-volume urea CUT&RUN protocol (Lov-U CUT&RUN), which has been shown to improve detection of transcriptional cofactors associated with chromatin-bound transcription factors[48]. Using isolated nuclei from 7-day-old adult W1118 flies, we compared IgG negative-control samples to samples incubated with an Atg8a antibody previously validated for both *Drosophila* Atg8 and human LC3/GABARAP detection (**Fig. 1A**).

**Figure 1.**
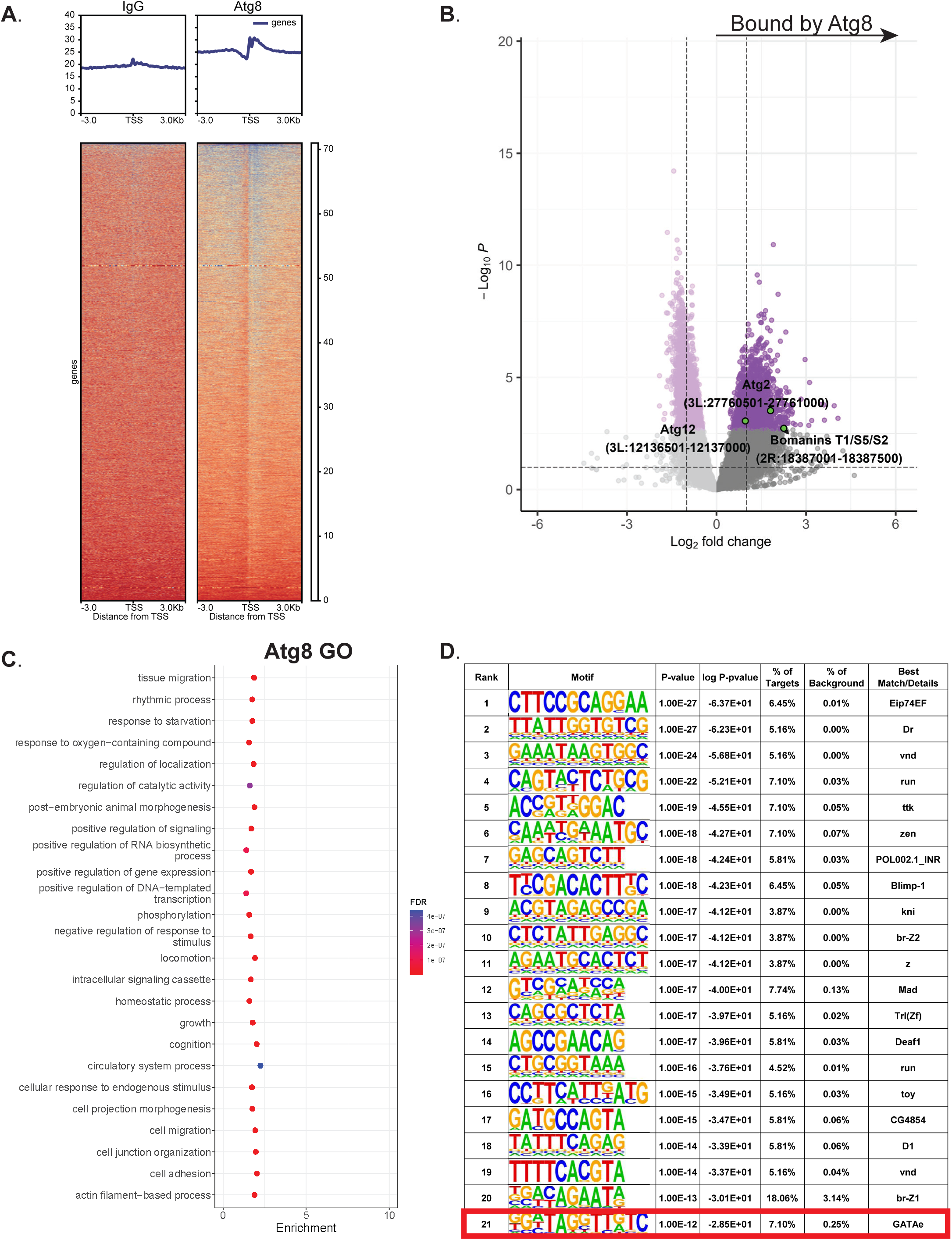
Atg8 Chromatin binding at active transcription sites in the basal state. A) overview of Atg8 chromatin binding in *W1118* 7-day-old adult flies on ND aligned to 5’ transcription start sites (TSS), compared to IgG negative control. The heatmap depicts the number of sequences relative to other TSS in arbitrary units with each row being a gene. Top graphs plot the mean occupancy across all TSS within +/- 3kb. B) Volcano plot of Atg8 CUT&RUN binding at each 500bp region of the Drosophila genome compared to IgG control. Positive log fold change indicates an Atg8-enriched chromatin region compared to IgG. Regions in purple indicate significantly enriched sites (padjust <0.05) compared to the control based on differential expression. Highlighted sites in green are selected regions found to be significantly enriched in Atg8 and identified as significantly enriched in SEACR analysis. C) Gene Ontology analysis of genes associated with SEACR significantly enriched sites of Atg8 binding. D) HOMER motif analysis of SEACR significantly enriched sites of Atg8 binding, with GATAe motif highlighted.

Heatmap analysis revealed a pronounced enrichment of Atg8 occupancy near transcription start sites across the *Drosophila* genome (**Fig. 1A–B**). Notably, this enrichment pattern resembled the localization profile of H2A.Z-containing nucleosomes, which are characteristically enriched at transcription start sites[49]. These findings suggest that Atg8 may function more broadly as a chromatin-associated[49] regulatory factor, consistent with prior reports implicating Atg8 in transcriptional control.

To identify candidate transcriptional regulators that may interact with Atg8, endogenous Atg8 was immunoprecipitated from isolated adult fat bodies using a Atg8a antibody, followed by immunoprecipitation mass spectrometry (IP-MS) analysis (**Supplemental Fig. 1B**). Among the proteins enriched in Atg8 pulldowns were several metabolic and autophagy-associated factors, including AGBE, Pgm1, Gyg, and Gys, which are involved in glycogen metabolism and pyruvate-associated pathways. Additional enriched proteins included Hsp60A, Hsp60B, Hsp60C, and vip2, which are associated with mitochondrial stress and apoptosis pathways (**Supplemental Fig. 1B**).

In addition to metabolic regulators, Atg8-associated complexes showed enrichment for proteins involved in heterochromatin formation and regulation of DNA-templated transcription. These included Histone 2A and HDAC1, both of which have previously been implicated in Atg8 acetylation[50]. Importantly, Histone 2A and HDAC1 are also core regulators of chromatin accessibility and transcriptional regulation, suggesting that Atg8 may participate in transcriptional control more broadly than previously appreciated[51, 52].

Comparison of Atg8-enriched chromatin regions against IgG controls identified significant enrichment at several autophagy-related loci, including Atg2 and Atg12, consistent with previous reports showing that Atg8 regulates autophagy-associated genes through interactions with Sequoia[26]. However, Atg8 occupancy was also detected at numerous genomic regions unrelated to canonical autophagy pathways (**Fig. 1B**). To further characterize Atg8-associated loci, we performed Sparse Enrichment Analysis for CUT&RUN (SEACR) and conducted Gene Ontology analysis on genes associated with significantly enriched peaks (**Fig. 1C**). This analysis identified Atg8 occupancy at genes involved in starvation response, metabolic homeostasis, and growth regulation. Notably, Atg8 binding was also enriched at genes associated with signaling pathways, transcriptional regulation, and cellular responses to endogenous stimuli, further supporting a broader role for Atg8 in homeostatic gene regulation.

Because Atg8 lacks intrinsic DNA-binding capability, its transcriptional regulatory activity likely depends on interactions with chromatin-associated cofactors or transcription factors. To identify candidate transcription factors associated with Atg8-bound regions, we performed motif enrichment analysis using the HOMER platform (**Fig. 1D**). Among the enriched motifs identified was the GATAe motif, a zinc-finger transcription factor motif associated with Notch signaling, midgut development, and κB-responsive promoters such as CecA1[53, 54]. Consistent with this observation, Atg8-enriched chromatin regions also included loci encoding several Bomanin antimicrobial peptides, including BomT1, BomS5, and BomS2, which are induced during gram-positive bacterial infection and are regulated by Dif-dependent κB signaling pathways[55, 56]. Together, these findings confirm Atg8 association with established autophagy-related targets while also demonstrating widespread chromatin occupancy at transcriptionally active regions associated with immune and transcriptional regulatory pathways[26].

### Atg8 associates with Dif. Nuclear localized under HSD and Immune stress response

Given that Atg8 functions both as a key mediator of extracellular signaling in addition to its canonical role in autophagy[13] and that its localization between the nucleus, cytoplasm, and extracellular space is tightly regulated in response to cellular stress and starvation[27], we next investigated whether obesogenic stress alters Atg8 abundance and cellular localization. To address this, we measured Atg8 levels in cellular lysates from flies maintained on either a normal diet (ND) or a high-sugar diet (HSD), which contains 30% more sugar than ND and has previously been shown to function as an obesogenic stressor[34].

At early stages of HSD exposure (3 days), Atg8 levels were significantly reduced in whole-cell lysates compared to ND controls (**Fig. 2A**). However, following chronic HSD exposure (14 days), a condition our work has previously associated with metabolic dysfunction, Atg8 accumulated significantly within the cellular lysate relative to ND controls[34, 57, 58]. To determine whether these changes were accompanied by altered subcellular localization, we performed immunohistochemistry on adult fat bodies using an anti-Atg8a antibody. In addition to increased overall Atg8 abundance, HSD-treated flies exhibited a marked increase in nuclear-localized Atg8 compared to ND controls (**Fig. 2C**).

**Figure 2.**
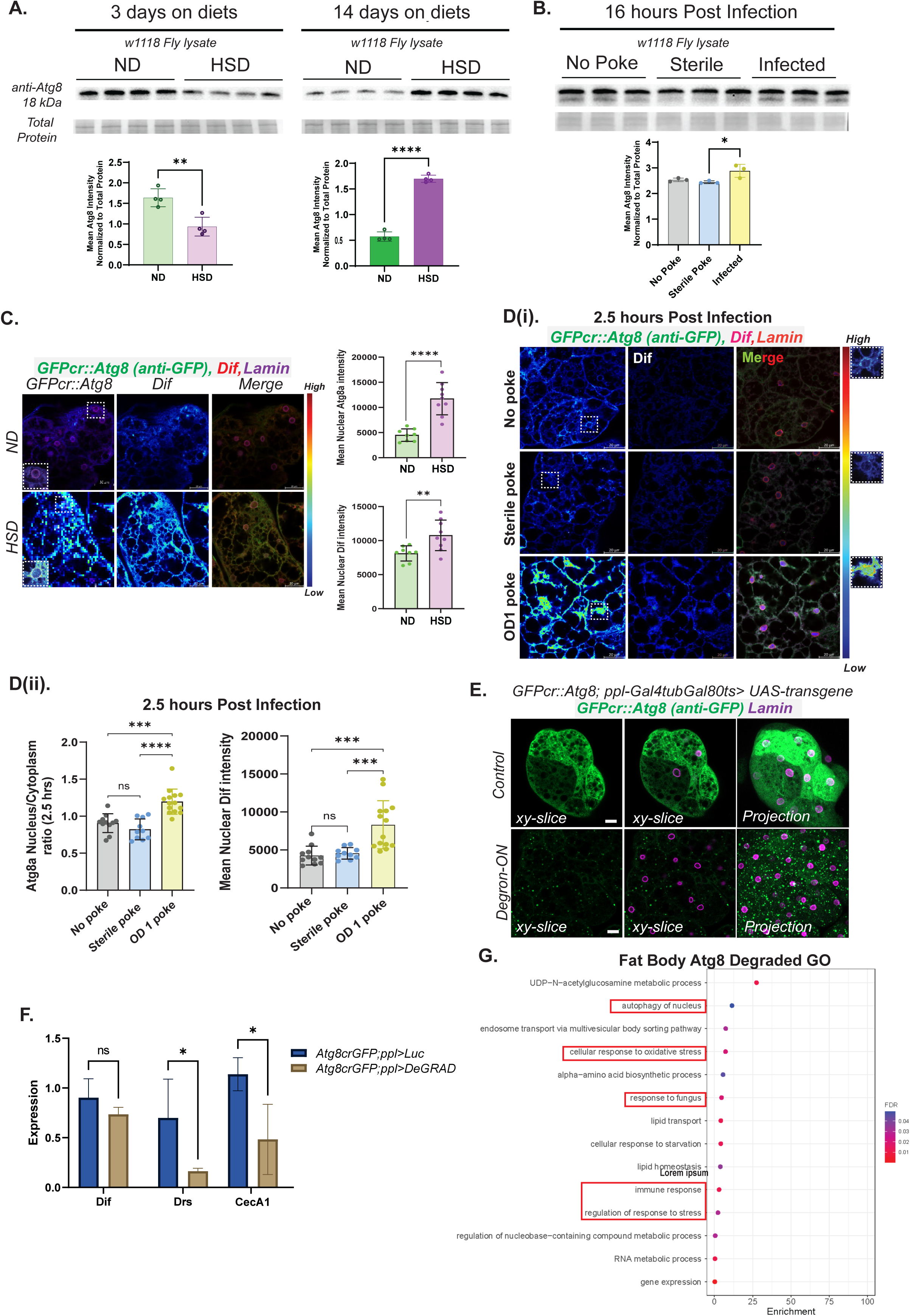
Atg8 associates with Dif. Nuclear localized under HSD and Immune stress response. A) Western blots of cellular lysate of *W1118* flies after 3 (left) or 14 (right) days on ND or HSD stained for Atg8 with quantification (bottom). B) Western blot stained for Atg8 in *W1118* flies infected with *E. Faecalis* 16 hours post infection, compared to controls with quantification (bottom). C) Atg8 localization in *W1118* fly fat bodies after 3 day on ND (top) or HSD (bottom), with Atg8 labeled in blue (left), nuclear lamin in pink (middle), and a merge (right). Quantification of mean nuclear intensity of Atg8 and Dif for each condition are shown (right bar graphs). D) Depicts Atg8 (left,blue), Dif (middle, purple) and merged (right) of *W1118* fly fat bodies 2.5 hours after no poke control (top) or injected with LB broth (middle), and *E. Faecalis* (bottom). The nucleus is labeled with lamin antibody (red). Mean nuclear intensity is of Atg8 and Dif is quantified (left, bar graphs) E) Degron-mediated knockdown reduces *GFP::Atg8* signal in the fat body. Representative confocal micrographs of fat body tissue from *GFP::Atg8; ppl-Gal4, tub-Gal80ts* flies expressing either *UAS-Luciferase* as a control (top) or *UAS-DegRAD-FP* (bottom) to induce degradation of GFP-tagged Atg8. Flies eclosed at 18°C, were shifted to the restrictive temperature, 29°C, for 2 weeks to activate transgene expression. *GFP::Atg8* was detected by anti-GFP staining, and Lamin marks nuclei. In control tissue, Atg8 shows broad nucleo-cytosolic localization with abundant GFP signal. In *UAS-DegRAD-FP*-expressing tissue, Atg8 signal is markedly reduced, with the remaining GFP-positive signal appearing as puncta, consistent with localization to possible degradative compartments. Single xy-slices are shown in the left and middle panels, and maximum-intensity projections of the entire tissue are shown in the right panels. F) qPCR validation confirmed that Dif-dependent AMPs, *Drosomycin* (Drs) and *Cecropin* (CecA1), were significantly downregulated following Atg8 degradation. G) Bulk RNA-seq of Atg8 degradation in the fat body under starvation conditions compared to Luciferase control and significant genes from Differential expression analysis (padj<0.05) was assessed for GO enrichment 30 fat bodies per sample, 3 biological replicates). qPCR of Scale bar, 10 μm. Blots are quantified with ImageJ and normalized to total protein. Statistical significance was assessed by t-test. error bars indicate standard deviation. *p<0.5,**p<0.1,***p<0.01,****p<0.001

These findings indicate that Atg8 localization is dynamically responsive to dietary stress and suggest that obesogenic conditions promote nuclear accumulation of Atg8, potentially enabling transcriptional regulatory activity. This observation is notable because canonical autophagic functions of Atg8 are typically associated with cytoplasmic localization. Importantly, HSD increased total cellular Atg8 levels globally, suggesting that dietary stress enhances Atg8 abundance in both nuclear and cytoplasmic compartments, potentially supporting simultaneous transcriptional regulation and autophagic homeostasis. Although these data suggested a role for Atg8 in stress-responsive nuclear processes, they did not directly establish chromatin association or transcriptional function.

Because our CUT&RUN analysis identified enrichment of Atg8 at immune-associated κB motifs, we next asked whether Atg8 nuclear localization also responds to immune stress. To test this, we utilized two Gram-positive bacteria, *Enterococcus faecalis* and *Lysinibacillus fusiformis*, both having well-documented and strong responses to the Toll pathway, eliciting Dif-dependent AMP production[59, 60]. *W1118* flies were infected with *E. faecalis* and Atg8 levels in fat body lysates were measured 16 hours post-infection (**Fig. 2B**). Infected flies showed a significant increase in cellular Atg8 compared to both unmanipulated and injection-control flies, demonstrating that infection itself, rather than injury, induces Atg8 accumulation. Immunohistochemistry was performed 2.5 hours following *E. faecalis* infection revealed a global increase in Atg8 abundance, along with significantly enhanced nuclear localization compared to controls (**Fig. 2D**).

Given the observed enrichment of Atg8 at κB-associated chromatin regions, we additionally examined the localization of the endogenous NF-κB transcription factor Dif during infection. As expected, Dif exhibited robust nuclear accumulation in response to immune challenge and showed significant colocalization with Atg8 within the nucleus following infection (**Fig. 2D**). In contrast, Dif localization was unchanged in fat bodies from flies exposed to ND or HSD for 3 days, and little Atg8–Dif colocalization was observed under these dietary conditions. These findings suggest that Atg8 responds broadly to multiple forms of cellular stress, whereas Dif activation is more specifically linked to immune challenge. Furthermore, the data indicate that Atg8 nuclear localization can occur independently of Dif signaling.

If Atg8 functions as a regulator of transcription, acute loss of Atg8 would be expected to produce distinct gene expression changes in pathways dependent on Atg8-mediated regulation. However, because Atg8 is essential for development and viability, traditional knockout approaches are difficult to interpret. To overcome this limitation, we developed a strategy for the inducible degradation of endogenous Atg8 specifically within the adult fat body. Using CRISPR, the endogenous Atg8 locus was N-terminally tagged with GFP to generate the *GFPcrAtg8* line. Homozygous *GFPcrAtg8* flies remained viable and fertile, indicating that the tagged protein retained functionality. Atg8 was then acutely degraded in adult fat bodies using a GFP-targeting degron system activated 7 days after eclosion. Validation experiments demonstrated a strong reduction in GFP signal in degron-expressing flies relative to luciferase controls under permissive activation conditions, confirming efficient depletion of endogenous Atg8 while maintaining fly viability (**Fig. 2E**).

Using this system, we performed bulk RNA-seq under starvation conditions to identify transcriptional changes induced by acute Atg8 depletion in the adult fat body (30 fat bodies per sample; three biological replicates). Gene ontology analysis revealed enrichment of autophagy-related pathways as well as immune-associated genes, including antimicrobial peptides (AMPs), further supporting both the established role of Atg8 in autophagy and its newly identified association with immune regulatory pathways from our CUT&RUN analysis (**Fig. 2G**). We further validated these RNA-seq findings by qPCR and confirmed that the Dif-dependent antimicrobial peptide genes *Drosomycin* (Drs) and *Cecropin A1* (CecA1) were significantly downregulated following Atg8 degradation. Taken together, these data demonstrate that Atg8 undergoes stress-responsive nuclear localization in response to both immune and metabolic challenge, and that acute loss of Atg8 disrupts immune-associated transcriptional programs, supporting a previously unrecognized role for Atg8 in gene regulation.

### Atg8 interaction critical for Dif infection response

Given the chromatin profile of Atg8, suggesting an unknown role in Dif’s gene regulation, we sought to determine if Atg8 physically interacted with Dif. An analysis of Dif’s protein sequence identified two Atg8 interacting motifs (AIMs) contained in Dif’s Rel Homology domain (**Fig. 3A**), giving Atg8 the potential to bind to Dif. To determine the importance of Atg8 interaction with Dif, we developed two CRISPR lines endogenously expressing Dif with and HA or GFP tag. One line contains the wildtype Dif sequence, the second contains Dif with the two AIM motifs mutated with two point mutations at the W and L sites (mutated to Alanines), rendering them inert and unable to bind to Atg8. We first wanted to determine if the AIM motif is critical for Dif binding to its transcription factor sites. To address this, we performed our Lov-U CUT&RUN with a Dif antibody we generated on our *DifWT*, *DifAIM*, and *Dif(1)* flies, with *Dif(1)* being a previously reported point mutation of Dif that loses its ability to bind DNA. We find that Dif occupancy is highest in *DifWT* and similar to *W1118* while *DifAIM* showed lower overall occupancy at TSS, and *Dif(1)* showed little to no occupancy, similar to IgG negative controls (**Fig. 3B**). This showed that the point mutations impacting Atg8 interacting with Dif did cause Dif to bind DNA at lower levels. However, it should be noted that this could be either due to the lack of Atg8 as a cofactor or the impact of the point mutations on the Dif’s Rel homology domain.

**Figure 3.**
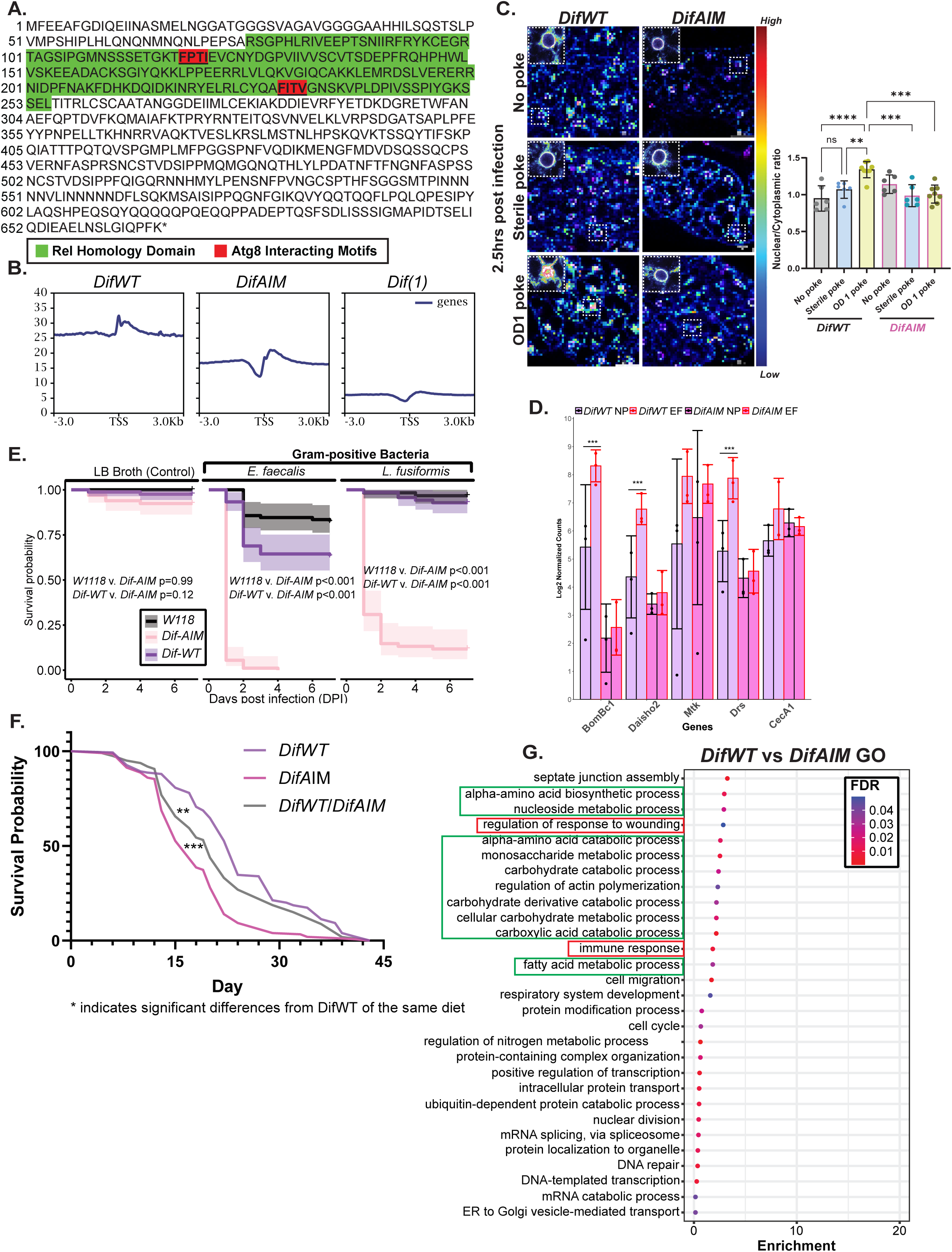
Atg8 interaction critical for Dif infection response. A) Dif protein sequence, green highlight details Rel Homology domain of Dif, red sites denote AIM sequences. Both AIM sites had double point mutations at the W and L sites of the AIM in *DifAIM* mutant flies. B) Dif Occupancy at TSS in *DifWT*, *DifAIM*, and *Dif(1)* flies. C) Nuclear localization of Dif in *DifWT* (left) and *DifAIM* (right) fly fat bodies after 2.5 hours E. Faecalis infection (bottom) compared to no poke control (top) and LB Broth injection (middle). D) Gene Expression of select downstream Dif targets in RNAseq analysis of uninfected (black outline) vs infected (red outline) in *DifWT* (purple) and *DifAIM* (pink) flies. E) Survival curves for *W1118*, *DifWT*, and *DifAIM* upon Gram-positive infection and F) under HSD. G) GO Enrichment of significantly differentially expressed genes in *DifWT* vs *DifAIM*. Error bars indicate standard deviation. Significance was determined by two-way ANOVA with Bonferroni post hoc correction. *p<0.5,**p<0.1,***p<0.01,****p<0.001

Having identified two putative AIMs within the Rel homology domain of Dif (**Fig. 3A**) and shown that mutating these residues reduces Dif chromatin occupancy at transcription start sites (**Fig. 3B**), we next asked whether the same AIM residues are required for Dif nuclear deployment during immune challenge. We examined endogenous Dif subcellular localization in adult fat bodies from *DifWT* and *DifAIM* flies under three conditions: unmanipulated control, sterile LB-broth injection (sterile poke), and *E. faecalis* infection (**Fig. 3C**). Consistent with the well-established Toll-pathway response, *DifWT* translocated to the nucleus between 30 minutes and 2.5 hours after *E. faecalis* infection, with the nuclear-to-cytoplasmic ratio of Dif rising significantly only under productive infection (**Fig. 3C**, N/C ratio quantification). Sterile poke did not elicit comparable nuclear accumulation, confirming that translocation is infection-driven rather than a generalized wound response. *DifAIM,* despite being expressed at levels comparable to *DifWT*, failed to enter the nucleus on infection; the *DifAIM* signal is observed at the nuclear lamina, with no measurable rise in N/C ratio across any of the three conditions (**Fig. 3C**). Together, these data indicate that intact AIM residues are required for productive Dif nuclear import beyond the perinuclear compartment, and suggest a functional role for the putative Dif AIMs in the immune response. Consistent with this impaired nuclear deployment of *DifAIM*, RNA expression of direct Dif downstream targets, including *Drosomycin* (Drs) and *Cecropin A1* (CecA1), was significantly lower in *DifAIM* than in *DifWT* flies (**Supplemental Fig. 3B**), indicating that intact Dif AIMs are required for Dif-driven antimicrobial peptide induction.

Given that Dif is critical in its response to Gram-positive infection, we next compared RNAseq gene expression profiles of our *DifWT* flies compared to *DifAIM* under uninfected and *E. Faecalis* infection conditions, 6 hours post infection (**Fig. 3D**). We found that when looking at *Bomanins* and *Daishos* expression, direct downstream targets of Dif[55, 61], we found that upon infection, *DifWT* flies showed a robust and significant increase in expression of these antimicrobial peptides on infection when compared to uninfected controls. Conversely, *DifAIM* flies show no significant change to any of these AMPs when infected, displaying an impaired gene response to infection for Dif downstream targets in *DifAIM* flies.

We posited that this lack of response to infection in AMP production would have physiological implications for the fly under infection. To determine this we performed survival curves of *W1118*, *DifWT*, and *DifAIM* flies to determine if they could survive a Gram-positive infection challenge. *W1118* flies showed high survival to both Gram-postive bacteria and *DifWT* showed comparable survival with 75% or more surviving the Gram-positive bacteria challenge, although they displayed lower survival in the *E. Faecalis* infection (**Fig. 3E**). *DifAIM* flies, however, were very sensitive to Gram-positive infection with almost all flies dying after 2 day post infection. Our data indicates that the lack of interaction with Atg8 led to clear deficiencies in Dif’s ability to appropriately regulate gene expression, which in turn led *DifAIM* flies to be highly sensitive to infection. Given Dif’s apparent association with Atg8, we wanted to know if *DifAIM* flies were also susceptible to other stressors such as dietary stress. To address this, we put *DifWT*, *DifAIM*, and *DifWT*/*DifAIM* heterozygotes on HSD and compared survival curves (**Fig. 3F**). Interestingly, *DifAIM* indeed showed significantly lower survival in *DifWT* with Heterozygotes having a reduced but still significant decrease in survival. Given our previous data that Dif is unassociated with Dietary stress response, we posit that this effect may be due to impaired Atg8 chromatin localization due to interactions with an impaired Dif, rather than due to changes in Dif gene regulation. We then performed RNAseq, comparing whole body *DifWT* flies to *DifAIM* flies (**Fig. 3G**). When looking at the significantly differentially expressed genes and assessing their gene ontology, we identified that not only were immune genes being impacted, but many metabolic processes. This indicated that flies with impaired Dif-Atg8 interaction showed impacts to many biological processes, including those outside of Dif’s canonical immune function. Taken together, these findings demonstrate that Atg8–Dif interaction is essential for proper Dif-dependent chromatin occupancy, antimicrobial gene activation, and organismal survival during stress, revealing that Atg8 functions as a critical co-regulator of immune and metabolic transcriptional programs beyond its canonical role in autophagy.

### Atg8 displays distinct regulation of Immune genes during infection and signaling during HSD stress

Having established that Atg8 associates with chromatin in response to distinct stress conditions, we next sought to determine which transcriptional programs are regulated by Atg8 during these stress responses and whether Atg8-dependent regulation differs depending on the type of stress encountered. To address this, we performed Lov-U CUT&RUN on *DifWT* flies under uninfected conditions and 6 hours following Enterococcus faecalis infection. For an unbiased comparison of chromatin occupancy, the entire *Drosophila melanogaster* genome was segmented into 500 bp regions and analyzed for differential Atg8 binding between infected and uninfected flies (**Fig. 4A**). In parallel, RNA-seq analysis was performed on uninfected and infected *DifWT* flies (**Supplemental Fig. 4B**) to identify genes that exhibited both significant Atg8 enrichment by SEACR analysis and increased transcriptional expression during infection.

**Figure 4.**
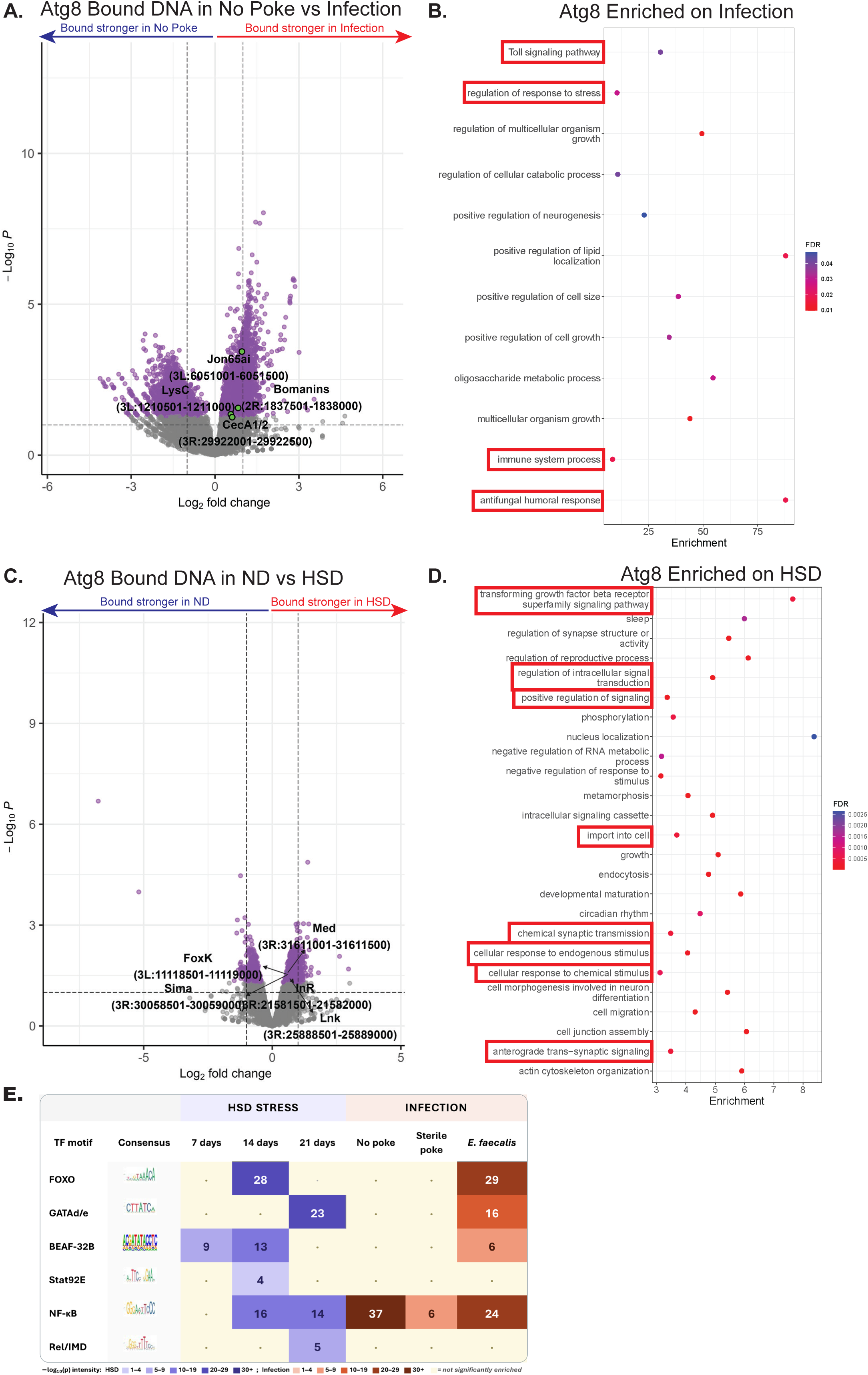
Atg8 displays distinct regulation of Immune genes during infection and signaling during HSD stress. A) Volcano plot comparing Atg8 chromatin binding between uninfected and 6 hr infected flies in *DifWT* and corresponding B) Gene ontology of genes upregulated on infection in RNAseq analysis (supplemental figure 4B) and significantly increased occupancy by Atg8 by SEACR analysis. Genes highlighted in green are selected AMP and immune factors impacted by Atg8 occupancy. Infection leads to significantly increased binding of Atg8 at AMP sites and immune genes (red highlights). C) Volcano plot comparing Atg8 chromatin binding between *W1118* on ND vs HSD for 14 days, and corresponding D) Gene ontology of genes upregulated on infection in RNAseq analysis (supplemental figure 4A) and significantly increased occupancy by Atg8 by SEACR analysis. Genes highlighted in green are select metabolic and signaling factors impacted by Atg8 occupancy. HSD leads to significantly increased binding of Atg8 at genes required for signaling (Red highlights). E) Summary table of Motif analysis using HOMER of chromatin regions bound by Atg8 using SEACR based on days under HSD or infection conditions.

Among the genes significantly associated with Atg8 during infection were several key immune regulators previously linked to Dif signaling, including members of the *Bomanin* family and *CecropinA1*. We additionally identified genes involved in gut immune homeostasis, including *Lysozyme C*, an antimicrobial peptide important for gut microbiota regulation[62], and Jon65Ai, a serine protease involved in intestinal homeostasis that is transcriptionally repressed during infection and starvation[63]. Gene ontology analysis of genes both significantly bound by Atg8 and transcriptionally upregulated during infection revealed strong enrichment for immune-associated pathways, including Toll signaling and antifungal humoral responses (**Fig. 4B**). Together, these findings demonstrate that during gram-positive bacterial infection, Atg8 becomes nuclear localized and selectively associates with immune-responsive loci involved in Toll pathway activation and antimicrobial peptide production.

We next examined whether Atg8 regulates distinct transcriptional programs during metabolic stress. Lov-U CUT&RUN was therefore performed on W1118 flies maintained for 14 days on either a normal diet (ND) or high-sugar diet (HSD). In parallel, RNA-seq analysis was conducted on corresponding ND and HSD samples. Using the same integrated analysis pipeline, we identified genes that showed significant Atg8 occupancy at promoter regions together with increased transcriptional expression under HSD conditions (**Fig. 4C**). Unlike the infection response, Atg8-associated genes under HSD were predominantly linked to metabolic and signaling pathways.

Notably, Atg8 occupancy was enriched at several genes involved in insulin signaling and metabolic regulation. These included FOXK, a major insulin-responsive transcription factor functionally opposed to FOXO and known to undergo nuclear localization following insulin stimulation[64, 65]; the *Insulin-like Receptor* (InR), a central regulator of *Drosophila* insulin-like peptide signaling required for growth and survival[66, 67]; and Lnk, an intracellular adaptor protein required for InR signaling[68]. In addition, Atg8 enrichment was identified at *Medea* (Med), a critical component of the SMAD/TGF-β signaling pathway involved in developmental patterning and transcriptional regulation[69]. Gene ontology analysis of Atg8-associated genes enriched during HSD exposure revealed pathways related to TGF-β signaling, intracellular import, and cellular responses to endogenous stimuli (**Fig. 4D**). These findings suggest that Atg8 regulation during dietary stress is mechanistically distinct from its role during infection and may instead function to prime intracellular signaling pathways involved in metabolic adaptation and homeostasis.

Because Atg8 exhibited distinct chromatin-binding profiles under infection and dietary stress, these findings suggested that Atg8 likely functions as a cofactor for multiple transcriptional regulators beyond the previously described Sequoia and Dif interactions. To further investigate this possibility, we performed HOMER motif enrichment analysis on all significantly enriched Atg8-binding regions identified by SEACR across multiple stress conditions, including 7-, 14-, and 21-day HSD exposure as well as no-poke, LB injection, and *E. faecalis* infection conditions (**Fig. 4E**).

This analysis revealed strong enrichment of κB-associated motifs during infection, with lower but detectable enrichment during prolonged obesogenic stress. Under both dietary and infection stress conditions, Atg8-associated regions were also enriched for FOXO and BEAF-32B motifs. BEAF-32B is an insulator-associated protein enriched near transcription start sites and broadly involved in chromatin organization and transcriptional regulation[70]. In contrast, Stat92E motifs, which regulate Jak–STAT signaling downstream of Unpaired ligands[71], were enriched during dietary stress but largely absent during infection. Interestingly, motifs associated with Rel/IMD signaling were notably absent under both HSD and infection conditions. This finding suggests that Atg8 does not function as a general immune-response regulator but instead preferentially associates with Toll pathway–related transcriptional programs during gram-positive infection. Furthermore, enrichment of Stat92E-associated motifs during HSD may reflect Atg8 involvement in metabolic signaling pathways linked to leptin-like Unpaired signaling[72].

Together, our findings establish Atg8 as a dynamic chromatin-associated regulator that links cellular stress responses to context-specific transcriptional programs in *Drosophila melanogaster*. Using Lov-U CUT&RUN, RNA-seq, motif enrichment analysis, and inducible genetic perturbation approaches, we demonstrate that Atg8 localizes to transcriptionally active chromatin regions, accumulates in the nucleus in response to both metabolic and immune stress, and regulates distinct gene networks depending on the physiological challenge encountered. Under gram-positive bacterial infection, Atg8 selectively associates with κB-responsive immune loci and is required for proper Dif-dependent antimicrobial peptide expression and host survival. In contrast, during chronic high-sugar diet stress, Atg8 preferentially associates with genes involved in metabolic signaling, insulin responsiveness, and cellular homeostasis. Mechanistically, we identify direct interaction between Atg8 and the NF-κB transcription factor Dif through conserved AIM motifs, revealing that disruption of this interaction impairs Dif chromatin occupancy, downstream immune gene activation, and organismal stress resistance. Collectively, these results expand the functional role of Atg8 beyond canonical autophagy and extracellular signaling, positioning Atg8 as a stress-responsive transcriptional cofactor that coordinates chromatin-associated regulatory programs involved in immunity, metabolism, and homeostatic adaptation.

## DISCUSSION

Here we report that endogenous Atg8 occupies promoter-proximal chromatin broadly across the adult *Drosophila* fat body genome, undergoes regulated nuclear accumulation in response to both gram-positive bacterial infection and chronic high-sugar diet (HSD), and engages the NF-κB transcription factor Dif through two conserved Atg8-interacting motifs (AIMs) within Dif’s Rel Homology domain. CRISPR-engineered AIM-mutant Dif retains expression but fails to translocate to the nucleus on infection, exhibits markedly impaired chromatin occupancy, and renders flies susceptible to both infection and chronic HSD. Comparative Atg8 chromatin profiling reveals partial overlap and substantial context-specific redeployment between these stressors. Together, these findings position Atg8 as a stress-responsive chromatin-associated cofactor, with NF-κB/Dif as one mechanistic client through which Atg8 exerts this role. Our results raise a set of open questions about how Atg8’s biochemistry, including its AIM/LIR partnership, its lipidation, its membrane-association capacity, and its association with histone-containing chromatin, supports this newly identified nuclear function. We discuss each in turn.

A first puzzle concerns the relationship between Dif and Atg8 chromatin occupancy in the AIM-mutant background. *DifAIM* and *Dif(1)*, a canonical DNA-binding-deficient allele[73], produce distinct chromatin profiles: *Dif(1)* shows essentially no occupancy at its target loci, as expected for a DNA-binding mutant, whereas *DifAIM* retains protein expression and intermediate residual chromatin occupancy at a subset of target loci. More strikingly, Atg8 chromatin engagement is disproportionately compromised in the *DifAIM* background relative to *Dif(1)*, even though *DifAIM* still nominally produces a Dif protein. One interpretation consistent with our nuclear localization data is that *DifAIM*, unlike *Dif(1)*, exposes a surface in the Rel Homology domain that *actively interferes* with Atg8’s translocation to the nucleus, sequestering Atg8 in the cytoplasm or otherwise preventing its productive nuclear entry. This model would assign Atg8 a dual role: first, to support the cytosol-to-nucleus shuttling of Dif itself (potentially as a chaperone or licensing partner during nuclear import), and second, to function as a chromatin-engagement cofactor once Dif occupies its κB sites. The dual-role model also accounts for the observation that AIM-mutant Dif itself fails to translocate to the nucleus on infection, despite intact expression. Whether Atg8 enters the nucleus with Dif, or whether Atg8 nuclear residency is regulated independently and Dif simply requires AIM-mediated docking on chromatin once nuclear, will be an important question for future work.

The susceptibility of *DifAIM* flies to chronic HSD raises a related interpretive question. Dif is classically considered an immune transcription factor, and Dif itself does not show dramatic nuclear translocation under our HSD condition. Why, then, do *DifAIM* flies fail to survive prolonged HSD? Two non-exclusive possibilities merit consideration. First, Dif may directly occupy a subset of HSD-responsive loci that have been underappreciated in the immune-centric literature on Toll/Dif function; AIM-mediated Atg8 partnership at these loci could be required for metabolic resilience even in the absence of overt nuclear translocation. Second, AIM-mutant Dif may act as a dominant *sink* for Atg8, interfering with Atg8’s chromatin function at the broader set of HSD-responsive metabolic loci (FOXK, *Stat92E*, insulin-pathway, and SMAD-linked motifs we identify) that depend on Atg8 but not on Dif. We currently favor a model that combines both: Dif and Atg8 form a regulatory partnership that, when disrupted by AIM mutation, both eliminates a focal κB-linked transcriptional output and compromises a broader Atg8-dependent chromatin program upon which metabolic homeostasis depends. Distinguishing direct Dif-at-HSD-loci effects from indirect Atg8-titration effects remains an open question for future work.

Beyond its AIM-mediated partnership with Dif, our data suggest that Atg8 engages chromatin through additional, partner-independent modes. The Atg8 CUT&RUN profile at baseline (**Fig. 1A**) shows a gene-end and TSS distribution that resembles homotypic H2A.v-containing nucleosomes more closely than a narrow sequence-specific transcription-factor footprint[49]. Our Atg8 fat-body immunoprecipitation-mass spectrometry similarly recovers H2A.v, HDAC1, HP1c, and Mi-2 among the enriched partners, a set of histone and chromatin-remodeling factors with no obvious AIM/LIR signature. Together, these observations suggest that Atg8 may associate with chromatin as a nucleosome-proximal factor in addition to, or independently of, sequence-specific transcription-factor partnerships.

Two mechanisms could account for this histone-associated engagement. The first is cofactor scaffolding, in which Atg8 is recruited indirectly to chromatin through histone-associated partner proteins. Atg8 acetylation is regulated by Sir2 and DOR[27] and by HDAC1[50], all of which are chromatin-associated factors that could, in principle, bridge Atg8 to histone-containing chromatin. Mi-2 and HP1c, additional chromatin-remodeling factors recovered in our IP-MS data, represent further candidate scaffolds. The second possibility is direct covalent modification of histone or histone-associated substrates by Atg8 itself. Histones are well-established substrates of ubiquitin-family modification: monoubiquitination of histone H2A and H2B at chromatin is a fundamental mode by which the ubiquitin system shapes transcriptional output[2–4], and the histone variant H2A.Z is itself ubiquitylated in mammalian cells, where this mark has been linked to both gene activation and gene silencing, depending on chromatin context[74]. Against this background, the concordance we observe between Atg8 chromatin occupancy and H2A.v-containing nucleosomes is particularly suggestive. Atg8/LC3 family proteins, in addition to their canonical conjugation to phosphatidylethanolamine, can themselves be covalently attached to other cellular proteins, including the Atg3 and Atg7 enzymes of their own conjugation machinery, in a process termed Atg8ylation[75–78]. Given Atg8’s structural relationship to ubiquitin and this established protein-substrate biochemistry, Atg8ylation of histone substrates emerges as a coherent candidate mechanism rather than a remote one. One particularly intriguing version of this model is that Atg8ylation functions as an *alternative* histone mark to ubiquitylation at the same lysine residue, analogous to the regulatory competition between methylation and acetylation at H3K27, producing distinct transcriptional outcomes depending on which modifier engages the substrate. ATG4 proteases, which reverse Atg8ylation[75], would provide the dynamic regulation expected of such a stress-responsive histone PTM.

Several candidate substrates remain to be distinguished. Atg8ylation could target H2A.v itself, the canonical H2A or H2B lysines that receive ubiquitin, or histone-associated cofactors. None has yet been directly tested. The question is increasingly tractable. Ordering Atg8 chromatin occupancy against established active-promoter marks such as H3K4me3 would clarify the chromatin context in which Atg8 engages. Mass-spectrometric analysis of purified Atg8 complexes would identify the modified residues directly. We highlight Atg8ylation as one of several candidate mechanisms, but a particularly plausible one, that the field will need to consider as it defines how the Atg8/LC3 system has been adapted for transcriptional regulation, alongside the partner-binding and acetylation modes already established.

A second, related question concerns the lipidation status of nuclear Atg8 and its relationship to the nuclear envelope. Atg8’s covalent conjugation to phosphatidylethanolamine (PE) is essential for autophagosomal membrane association, but whether nuclear Atg8 is similarly lipidated, or whether it operates as an unconjugated soluble cofactor, remains unresolved. The possibility that lipidated Atg8 engages the nuclear envelope draws support from prior work showing that LC3 binds Lamin B1 directly via an LIR-like motif in Lamin B1, recruiting Lamin B1 for autophagic degradation during oncogenic and senescence-associated stress[79]. By analogy, lipidated Atg8 at the inner nuclear membrane could organize histone-containing chromatin non-covalently from the lamina inward, providing a structural scaffold by which the nuclear envelope licenses stress-responsive chromatin engagement. An alternative model is that nuclear Atg8 is largely unconjugated and acts through soluble cofactor interactions independent of membrane attachment. Distinguishing these possibilities is an important direction for future work.

A striking pattern emerges from comparative motif analysis of Atg8-occupied chromatin across infection and HSD. FOXO and BEAF-32B motifs are co-enriched at Atg8 peaks uniquely under both stressors, two physiologically orthogonal challenges, one chronic and metabolic, the other acute and immune. The Mad/Med BMP-SMAD motif also recurs across both conditions. Three shared motif hits across independent stressors point to a stress-responsive chromatin signature that is not strictly immune and not strictly metabolic, but instead a more general state-responsive grammar.

The convergence at FOXO motifs in particular invites a more explicit interpretation. FOXO occupies an unusual position in adipose biology, sitting at the intersection of two regulatory programs that both involve the Atg8/LC3 system: autophagy gene induction during nutrient deprivation, and innate immune homeostasis through tonic antimicrobial peptide expression in *Drosophila* fat body[80]. Under acute starvation or infection, nuclear translocation of FOXO drives an adaptive transcriptional program that engages both arms: turnover of intracellular components through autophagy and humoral defense through AMP induction. We have previously shown that under chronic high-sugar diet, however, nuclear FOXO accumulates by a distinct mechanism. At 14 days of HSD, fat-body insulin resistance disrupts the insulin/Akt signaling that normally retains FOXO in the cytoplasm, producing persistent nuclear FOXO in the absence of an actual nutrient deficit[34]. The novel finding here is therefore not nuclear FOXO itself, but the physical co-occupancy of Atg8 at FOXO target promoters at this same insulin-resistant timepoint, suggesting that Atg8 acts as a transcriptional cofactor at FOXO targets during chronic metabolic dysfunction.

This convergence is mechanistically suggestive: a single conserved molecular handle (Atg8) co-occupies the chromatin of a transcription factor (FOXO) that regulates both autophagy and immune programs, in a physiological state (insulin resistance) where both programs are chronically engaged. Whether Atg8–FOXO co-occupancy under HSD supports adaptive responses to dietary stress or instead sustains a *maladaptive* chronic-stress program is a question this study cannot yet resolve. Signaling machinery initially evolved for acute adaptive responses often becomes harmful when chronically engaged, a defining feature of chronic low-grade inflammation in obesity[81]. Atg8 deployment at FOXO targets during sustained nutrient surplus may be one such case. Distinguishing adaptive from maladaptive Atg8–FOXO engagement, and determining whether the same molecular handle that confers infection resilience under acute challenge contributes to metabolic dysfunction when chronically deployed, will be an important goal for future work.

Layered on this shared FOXO/BEAF-32B/Mad backbone, each stressor recruits Atg8 to additional, condition-specific partner motifs: *tin*/NKX2.5 and GATAd under acute infection; FOXK, *Stat92E*, insulin-pathway, and srp/GATA family under HSD. Curiously, *dl* (Dorsal/Toll NF-κB) appears at Atg8-occupied promoters in sterile-wound controls but is absent under productive *E. faecalis* infection, suggesting that Atg8 redirects away from generic Toll activation and toward Dif-specific outputs during true pathogen challenge. We propose that Atg8 deploys at a shared chromatin core defined by FOXO/BEAF-32B/Mad grammar, with additional context-specific motifs recruited by the upstream physiological state.

Our findings open more questions than they close, and we view them as the starting point of a framework rather than a complete model. Several mechanistic questions remain: whether Atg8 acts upstream of Dif at the level of nuclear import, whether Atg8 chromatin engagement requires its PE-lipidation status, whether Atg8 contacts histones through partner-protein scaffolding, Atg8ylation of histone or cofactor substrates, or other modes of histone-proximal association, and whether Lamin-mediated INM tethering supports Atg8’s broader histone-associated chromatin landscape. Each of these questions opens a direction for future investigation. An important question for the field will be whether the chromatin-engagement principles defined here in adult *Drosophila* fat body generalize to mammalian adipose biology, including the LC3 and GABARAP family members in primary human adipocytes under conditions of metabolic and immune stress that converge in adipose tissue. We posit that the Atg8/LC3 family, long understood as the central effector of macroautophagy, should also be considered a stress-responsive cofactor in transcriptional regulation, with the AIM/LIR motif as one demonstrated interface among several mechanisms (including Atg8ylation of protein substrates) by which the autophagy machinery has been repurposed to license stress-responsive chromatin. Defining the full scope of these mechanisms will inform how stressed adipose tissue, and adipose-derived signals more broadly, integrates immune and metabolic information at the level of gene expression, with relevance to therapeutic strategies in chronic inflammation, obesity, and the metabolic-immune diseases that connect them.

## ACKNOWLEDGEMENTS

We thank Steven Henikoff for advice on CUT&RUN modifications and for insightful discussions on chromatin biology. We are grateful to Dominique Ferrandon and Ylva Engström for generously sharing Dif antibodies. We thank Terry L. Hafer, whose initial work identified the AIM/LIR motifs in Dif. Jordan Wong provided technical support for HSD survival curves, and Merriam Al-Maiahi provided technical support for immunohistochemistry, dissections, and fly husbandry. We thank the Fred Hutchinson Cancer Center Shared Resources for technical support, including the Genomics & Bioinformatics Shared Resource, especially Alex Zevin, the Proteomics & Metabolomics Shared Resource, and the Cellular Imaging Shared Resource, with particular thanks to Dr. Julien Dubrulle for morphometric analyses. We thank members of the Rajan and Unckless laboratories for helpful discussions and feedback. The graphical abstract was created in BioRender (biorender.com) under the Fred Hutchinson Cancer Center institutional license. Stocks obtained from the Bloomington Drosophila Stock Center (NIH P40OD018537) were used in this study.

This work was supported by the National Institute of General Medical Sciences of the National Institutes of Health under award R35GM124593 to A.R., and by NSF DEB grant 2330095 to R.L.U. This research was also supported by NIH P30 CA015704 of the Fred Hutch/University of Washington/Seattle Children’s Cancer Consortium, which includes the Genomics & Bioinformatics Shared Resource (RRID:SCR_022606), the Cellular Imaging Shared Resource (RRID:SCR_022609), and the Proteomics & Metabolomics Shared Resource (RRID:SCR_022618).

## METHODS

### Animals used and rearing conditions

The following fly strains were used in this study: *W1118*, *DifcrWT*, *DifcrAIM*, *Dif(1)*, *GFP::Atg8*, *UAS-Luciferase*, *UAS-DeGradFP*, *ppLGAL4,tubGAL80ts*, *GFP::Atg8;ppL,tubGAL80ts>UAS-Luc*, and *Atg8;deGradFP;ppL,tubGAL80ts>UAS-Degrad*. Sequence analysis of Dif identified two putative Atg8-interacting motifs (AIMs; red) within the DNA-binding domain (green). Using CRISPR, we generated an endogenous C-terminally HA-tagged Dif line (*DifWT*) and a mutant line (*DifAIM*) in which the first and fourth residues of both AIMs were mutated to alanine to disrupt Atg8 interaction. Both lines were homozygous viable and fertile.

To enable acute degradation of Atg8 in the adult fat body, we generated a *GFP::Atg8* line by N-terminally tagging the endogenous Atg8 locus with GFP using CRISPR. Homozygous *GFP::Atg8* flies were viable and fertile, indicating that the GFP-tagged protein remained functional. Acute degradation of Atg8 in adult fat bodies was achieved by crossing *GFP::Atg8;ppL-Gal4,tubGAL80ts* flies with either *UAS-DegradFP* or *UAS-Luciferase* control flies. For Atg8 degron experiments, flies were reared at 18°C after eclosion and then shifted to the restrictive temperature (29°C) for 2 weeks to induce transgene expression. All other flies were maintained at 25°C.

Only adult male flies were used in all experiments. Flies were sexed upon eclosion using brief CO2 anesthesia, which was not used again for the remainder of the experiment. Male flies were maintained on a normal diet (ND) for 7 days. The ND consisted of 15 g yeast, 8.6 g soy flour, 63 g corn flour, 5 g agar, 5 g malt, and 74 mL corn syrup per liter. After 7 days, flies were either maintained on ND or transferred to a high-sugar diet (HSD), which had the same base composition as ND, supplemented with an additional 300 g of sucrose per liter (30% increase). Flies remained on the assigned diet for the durations indicated in the figures (3 or 14 days).

### Infection Studies

Several days prior to infection, bacteria were streaked onto standard LB agar plates without antibiotics to generate isolated colonies. Two days before infection, individual colonies were picked using a sterile toothpick and inoculated into 2 mL TH broth for overnight incubation in a 37°C shaking incubator. On the day of infection, liquid cultures were serially diluted and adjusted by spectrophotometry to an optical density (OD600) of 1.0 for both Enterococcus faecalis and Lactobacillus fusiformis.

Seven-day-old flies were anesthetized using either CO2 or cold shock and assigned to one of four treatment groups: no injection (No Poke), injection with sterile LB broth (LB control), injection with OD1 *E. faecalis*, or injection with OD1 *L. fusiformis*. Injections were performed below the wing hinge using a sterilized needle. Flies were anesthetized and injected in batches of 10, with all injections completed within 1 hour. The injection needle was sterilized with 95% ethanol between treatment groups.

### ND/HSD Atg8a Western Blots

Adult male *Drosophila melanogaster* (*w1118*) were aged to 7 days and then maintained on either a normal diet (ND) or a high-sugar diet (HSD) for 3, 7, 14, or 21 days. For each condition, 90 flies were collected. Ten flies were reserved for whole-fly lysate preparation, while the remaining 80 flies were used for hemolymph collection. Whole-fly lysates were prepared by homogenizing 10 flies in ice-cold lysis buffer supplemented with HALT protease inhibitor using bead homogenization. Lysates were incubated on a rocker for 60 minutes at 4°C and then centrifuged at 10,000 × g for 5 minutes at 4°C to pellet debris. Protein concentrations were subsequently quantified using a BCA assay. For western blot analysis, whole-fly lysates were mixed with 4× Laemmli Sample Buffer, TCEP, and water, then boiled at 95°C for 5 minutes. Five micrograms of protein per sample were loaded onto 8–16% precast gels, with four biological replicates analyzed per condition. Proteins were transferred onto nitrocellulose membranes, which were then blocked for 1 hour in a 1:4 dilution of StartingBlock Blocking Buffer in TBS-T. Membranes were incubated overnight at 4°C with anti-Atg8a primary antibody (1:2000) diluted in the same blocking solution. After washing with TBS-T, membranes were incubated for 1 hour at room temperature with HRP-conjugated rabbit secondary antibody and developed using ECL reagents.

### Western Blotting on Infected Flies

Adult male *Drosophila melanogaster* (*w1118*) were assigned to one of three treatment conditions: unmanipulated control (No Poke), puncture with a sterile needle dipped in sterile Todd Hewitt broth (sterile poke), or puncture with a needle dipped in an overnight culture of *Enterococcus faecalis*. Following treatment, flies were maintained at 25°C for the indicated incubation period.

Protein concentrations were determined using a BCA assay. Five micrograms of protein per sample were resolved on 8–16% precast gels and transferred onto nitrocellulose membranes. Membranes were blocked for 1 hour in a 1:4 dilution of StartingBlock Blocking Buffer in TBS-T, followed by overnight incubation at 4°C with anti-Atg8a (bcam Catalog # 109364 Rb-GABARAP [EPR 4805]) primary antibody (1:2000) diluted in the same blocking solution. The following day, membranes were washed with TBS-T, incubated for 1 hour at room temperature with HRP-conjugated rabbit secondary antibody, and developed using ECL reagents.

### GFP-tagged Western Blots

Fifteen to twenty whole *Drosophila melanogaster* flies were homogenized using bead disruption in ice-cold lysis buffer supplemented with HALT protease inhibitor. Lysates were incubated on a rocker at 4°C for 60 minutes and then centrifuged at 10,000 × g for 5 minutes at 4°C. The clarified supernatant was collected, and protein concentration was quantified using a BCA assay.

For each of the three genotypes, 400 µg of lysate was incubated with 25 µL GFP-conjugated magnetic agarose beads (GFP-Trap) and rotated at 4°C for 2 hours. More than 100 µg of whole-fly lysate was reserved as the total input fraction before incubation. Following incubation, magnetic separation was used to collect the beads, and the remaining supernatant was retained as the unbound (flow-through) fraction. Beads were washed three times for 5 minutes each in the wash buffer to remove non-specifically bound proteins. Bound proteins were then eluted by boiling the beads in 4× Laemmli Sample Buffer, TCEP, and water at 95°C for 5 minutes.

For western blot analysis, 10 µg of the input and flow-through fractions, along with 10 µL of the immunoprecipitated fraction, were resolved on 8–16% precast gels and transferred onto nitrocellulose membranes. Membranes were blocked for 1 hour in a 1:4 dilution of StartingBlock Blocking Buffer in TBS-T and incubated overnight at 4°C with primary antibodies against GFP (1:5000), Atg8a (1:2000), or Dif (1:1000), diluted in the same blocking solution. The following day, membranes were washed with TBS-T, incubated for 1 hour at room temperature with HRP-conjugated rabbit secondary antibody, and developed using ECL reagents to visualize protein bands.

### Western Blot Quantification and Analysis

Western blot quantification was performed using GraphPad Prism and ImageJ. Chemiluminescent membrane images were imported into ImageJ as TIF files, with bands displayed as white signal on a black background. Band intensity was quantified using the rectangular selection tool while maintaining a consistent region of interest size across all samples. Raw intensity measurements were exported from ImageJ into Excel for normalization. Total protein staining, obtained using Pierce reversible protein stain immediately after membrane transfer, was quantified using the same ImageJ workflow. Signal intensity for each protein of interest was normalized to the corresponding total protein signal within the same lane. Normalized values were then plotted and analyzed in GraphPad Prism.

### Immunostaining

Adult male *Drosophila melanogaster* flies aged 7–10 days were used for fat body dissections. Fat bodies were dissected in ice-cold phosphate-buffered saline (PBS) and immediately fixed in 4% formaldehyde in PBS for 20 minutes at room temperature with gentle agitation. Following fixation, tissues were briefly rinsed in PBS and washed sequentially in PBST (PBS containing 1.0% Triton X-100) for 5, 10, and 20 minutes.

Tissues were then blocked for 1 hour at room temperature in PBS containing 0.3% Triton X-100 and 5% normal donkey serum (NDS). After blocking, samples were incubated overnight at 4°C with primary antibodies diluted in PBS containing 0.3% Triton X-100 and 5% NDS.

The following day, tissues were washed extensively in PBS containing 0.3% Triton X-100 for 2 hours, consisting of four 30-minute washes. Samples were subsequently re-blocked for 30 minutes at room temperature in PBS containing 0.3% Triton X-100 and 5% NDS, followed by incubation with the appropriate secondary antibodies (1:500 dilution) and DAPI for 3 hours at room temperature.

After secondary antibody incubation, tissues were washed for 1 hour in PBS containing 0.3% Triton X-100, followed by an additional 3–4 washes of 5–10 minutes each. Samples were then mounted in antifade mounting medium (e.g., SlowFade). Images were acquired using the Zeiss LSM 800 confocal microscope system and analyzed as described above.

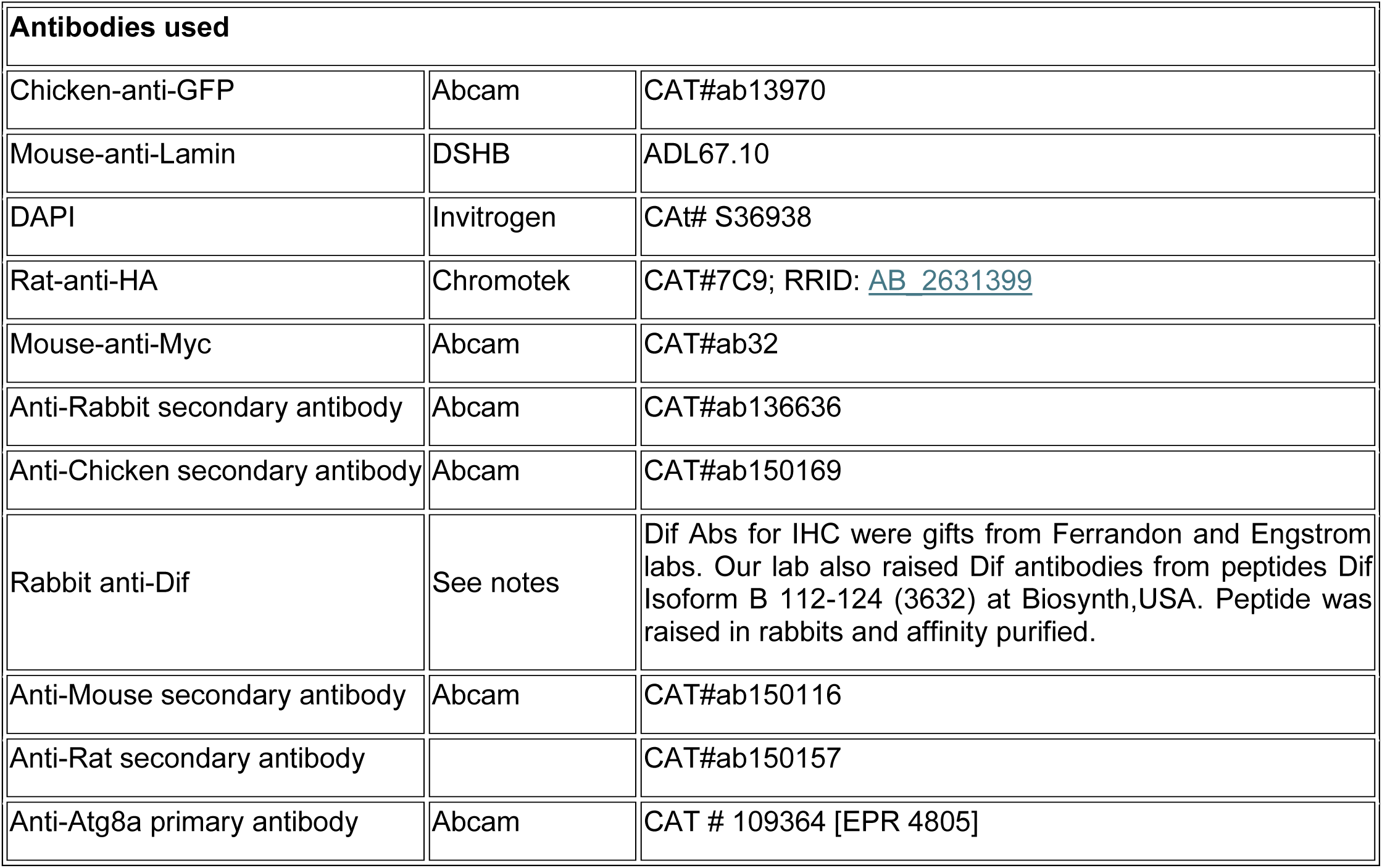

| Antibodies used |  |  |
| --- | --- | --- |
| Chicken-anti-GFP | Abcam | CAT#ab13970 |
| Mouse-anti-Lamin | DSHB | ADL67.10 |
| DAPI | Invitrogen | Cat# S36938 |
| Rat-anti-HA | Chromotek | CAT#7C9; RRID: AB_2631399 |
| Mouse-anti-Myc | Abcam | CAT#ab32 |
| Anti-Rabbit secondary antibody | Abcam | CAT#ab136636 |
| Anti-Chicken secondary antibody | Abcam | CAT#ab150169 |
| Rabbit anti-Dif | See notes | Dif Abs for IHC were gifts from Ferrandon and Engstrom labs. Our lab also raised Dif antibodies from peptides Dif Isoform B 112-124 (3632) at Biosynth,USA. Peptide was raised in rabbits and affinity purified. |
| Anti-Mouse secondary antibody | Abcam | CAT#ab150116 |
| Anti-Rat secondary antibody |  | CAT#ab150157 |
| Anti-Atg8a primary antibody | Abcam | CAT # 109364 [EPR 4805] |

### Adult *Drosophila* Fat body Confocal Image Analysis

A custom MATLAB analysis pipeline was developed to quantify Atg8 and Dif localization in fat body tissue from volumetric confocal imaging datasets. Confocal image files in CZI format were imported using the bfmatlab package, which provides Bio-Formats image reader and writer support for MATLAB. The lamin channel was used to segment nuclei in three dimensions through Otsu thresholding followed by standard morphological operations, including hole filling and size filtering. Fluorescence intensity measurements for the nucleus, cytoplasm, and nuclear envelope were performed on two-dimensional image planes to minimize signal variability caused by light scattering across tissue depth. For each nucleus, the focal plane with the largest nuclear cross-sectional area was selected for analysis. The nuclear envelope region was defined as the perimeter of the nuclear mask after thinning the mask by 6 pixels. To define the cytoplasmic region, the nuclear mask was expanded using a disk-shaped structural element with a 25-pixel radius, and the original nuclear area was excluded. This cytoplasmic mask was then intersected with a binarized image of the signal of interest to exclude lipid droplets from fluorescence quantification. The nucleo-cytoplasmic ratio was calculated as the mean fluorescence intensity within the nucleus divided by the mean fluorescence intensity within the cytoplasmic region for the signal of interest.

### Survival Assay

Survival assays were performed using male flies collected within a 24-hour eclosion window and aged for 7 days prior to experimentation. For high-sugar diet (HSD) survival assays, *DifWT*, *DifAIM*, and *DifWT*/*DifAIM* heterozygous flies were transferred to HSD media on Day 0 of the experiment, with 10 males housed per vial. Mortality was recorded daily until all flies had died. For infection survival assays, *W1118*, *DifWT*, and *DifAIM* flies were injected with either LB broth control or *Enterococcus faecalis*. Fly survival was monitored daily for 7 days or until all flies had died. All flies were maintained at 25°C under a 12-hour light/dark cycle throughout the duration of the experiments. Survival analyses were conducted using the Survival Curve module in GraphPad Prism, and statistical significance was determined using the Mantel–Cox (log-rank) test. More than 90 flies were analyzed per condition for each survival curve.

### RNA Sequencing

For each sample, 15 seven-day-old adult male *Drosophila melanogaster* flies were collected, with three biological replicates generated for each genotype and treatment condition.

For diet experiments, flies were maintained on either a normal diet (ND) or a high-sugar diet (HSD) for 14 days before flash freezing. For infection experiments, flies were infected with Enterococcus faecalis and compared to unmanipulated (No Poke) controls. Flies were flash frozen 6 hours post-infection.

For RNA extraction, zirconium beads and 30 µL TRI reagent (provided in the Direct-zol RNA Miniprep Kit) were added to each sample tube, and tissues were homogenized using a bullet blender. Total RNA was then isolated using the Zymo Research Direct-zol RNA Miniprep Kit (R2051-A). RNA quality and concentration were assessed using a SpectraMax i3 and an Agilent 4200 TapeStation.

RNA-seq libraries were prepared from total RNA using the TruSeq Stranded mRNA Kit. Library size distribution was validated using the Agilent 4200 TapeStation. Additional library quality control, indexed library pooling, and cluster optimization were performed using the Invitrogen Qubit 2.0 Fluorometer. Libraries were pooled in 35-plex and sequenced on an Illumina NextSeq P3 flow cell using paired-end 50 bp reads (PE50). We used FlyBase (release FB2026_01) to find information on phenotypes, function, and gene expression of significantly enriched genes[82].

### CUT&RUN

Unless otherwise specified, CUT&RUN experiments were performed using 7-day-old adult *Drosophila melanogaster* flies, with 10–15 whole flies used per biological replicate. A minimum of three biological replicates was included for each genotype, condition, and antibody treatment. Flies were flash-frozen in liquid nitrogen immediately after collection and stored at −80°C until processing.

Nuclei were isolated from whole-fly samples using the Rapid, Efficient, and Practical (REAP) subcellular fractionation method as previously described[83]. All procedures were performed on ice or in a 4°C cold room. Flies were homogenized using a bullet blender in ice-cold PBS supplemented with Roche cOmplete Protease Inhibitor Cocktail and Roche PhosSTOP at manufacturer-recommended concentrations. Large debris was removed by passing homogenates through a nylon mesh filter. Samples were briefly centrifuged, the supernatant was removed, and pellets were resuspended in 850 µL PBS containing 0.1% NP-40 and inhibitors. Following an additional centrifugation step, the supernatant was removed, leaving an enriched nuclear fraction for downstream processing. Each sample contained approximately 5 × 10^5^ nuclei.

CUT&RUN was performed using a modified protocol termed Lov-U CUT&RUN, which has been shown to improve detection of transcription factor and cofactor binding on chromatin[48]. The workflow followed the standard CUT&RUN procedure, including binding nuclei to Concanavalin A magnetic beads and overnight incubation at 4°C with the primary antibody of interest. Antibodies used in this study included IgG (negative control), H3K4me3 (positive control), endogenous GABARAP/Atg8, and Dif/NF-κB, each used at a 1:100 dilution. The Atg8a antibody was validated for use in both *Drosophila* and human adipocyte samples.

Following primary antibody incubation, samples were washed and incubated with EpiCypher Protein A/G–Micrococcal Nuclease fusion protein (pAG-MNase) for 30 minutes at 4°C to selectively target antibody-bound chromatin. Samples were then washed and incubated for 30 minutes with 100 mM calcium solution diluted 1:50 to activate MNase digestion. Unlike the standard CUT&RUN protocol, reactions were terminated using a low-volume stop buffer containing 8.8 M urea, 5 M NaCl, 0.5 M EDTA, 0.5 M EGTA, and 0.005% NP-40 for 1 hour. DNA was subsequently purified using Omega Bio-tek Mag-Bind TotalPure NGS Beads according to the manufacturer’s instructions. DNA concentration was quantified using Qubit fluorometric analysis.

NGS libraries were prepared using the CUTANA CUT&RUN Library Prep Kit (EpiCypher products 14-1001 and 14-1002) following the manufacturer’s protocol. Both primer sets supplied with the kit were used during library generation. Library concentration and quality were assessed using the Roche KAPA Library Quantification Kit. Samples were pooled at equimolar ratios into a final sequencing library with concentrations ranging from 7–14 nM. Primer dimers were removed using Omega Bio-tek Mag-Bind TotalPure NGS beads, and final pooled library quality was confirmed using a TapeStation. Sequencing was performed by the Fred Hutchinson Cancer Center Next Generation Sequencing Core using the Illumina NovaSeq X platform. Sequencing depth ranged from approximately 5–8 million reads per sample. Comparisons were performed within individual sequencing pools, with three biological replicates analyzed per antibody and experimental condition.

### CUT&RUN Analysis

Sequencing data for each sample were received in FASTA format and processed using a custom analysis pipeline based largely on the workflow described by Steven Henikoff and colleagues in their 2020 CUT&RUN analysis framework[84]. In brief, sequencing reads were aligned to the Dm6 *Drosophila melanogaster* genome using Bowtie2, SAMtools, and Picard. Aligned reads were processed into BAM files using BEDTools and SAMtools for downstream quantification and analysis.

Biological replicates were intersected using BEDTools, and enriched chromatin regions were identified using the Sparse Enrichment Analysis for CUT&RUN (SEACR) peak-calling algorithm. SEACR analysis was performed by comparing antibody-enriched samples against corresponding IgG negative-control datasets to identify statistically enriched binding regions for each protein of interest. Additional details regarding SEACR methodology are described in the original publication[85].

To identify enriched DNA-binding motifs within SEACR peaks, peak-associated sequences were analyzed using the HOMER motif discovery suite, specifically the findMotifs function, against known Drosophila motif databases. Motif enrichment significance was determined using HOMER-generated log p-values[86]. For differential chromatin occupancy analysis and volcano plot generation, the *Drosophila* genome was divided into unbiased 500 bp genomic bins. Read counts from aligned BAM files were quantified across these regions and compared between experimental conditions. Differential enrichment analysis was performed in R using the DESeq2 package with three biological replicates per condition. Corresponding volcano plots were generated to visualize significantly altered chromatin regions between treatments. Genes were classified as significantly regulated only if they met all of the following criteria: 1) The associated chromatin region showed significant differential enrichment in CUT&RUN volcano plot analysis (p < 0.05). 2) The same chromatin region was independently identified as significantly enriched by SEACR peak calling. 3) The corresponding gene, or its promoter region, exhibited significant differential expression in matched RNA-seq datasets (p < 0.05). We used FlyBase (release FB2026_01) to find information on chromosome location, phenotypes, function, and associated gene expression of significantly enriched chromatin regions[82].

---

**Figure 1 Supplement.**
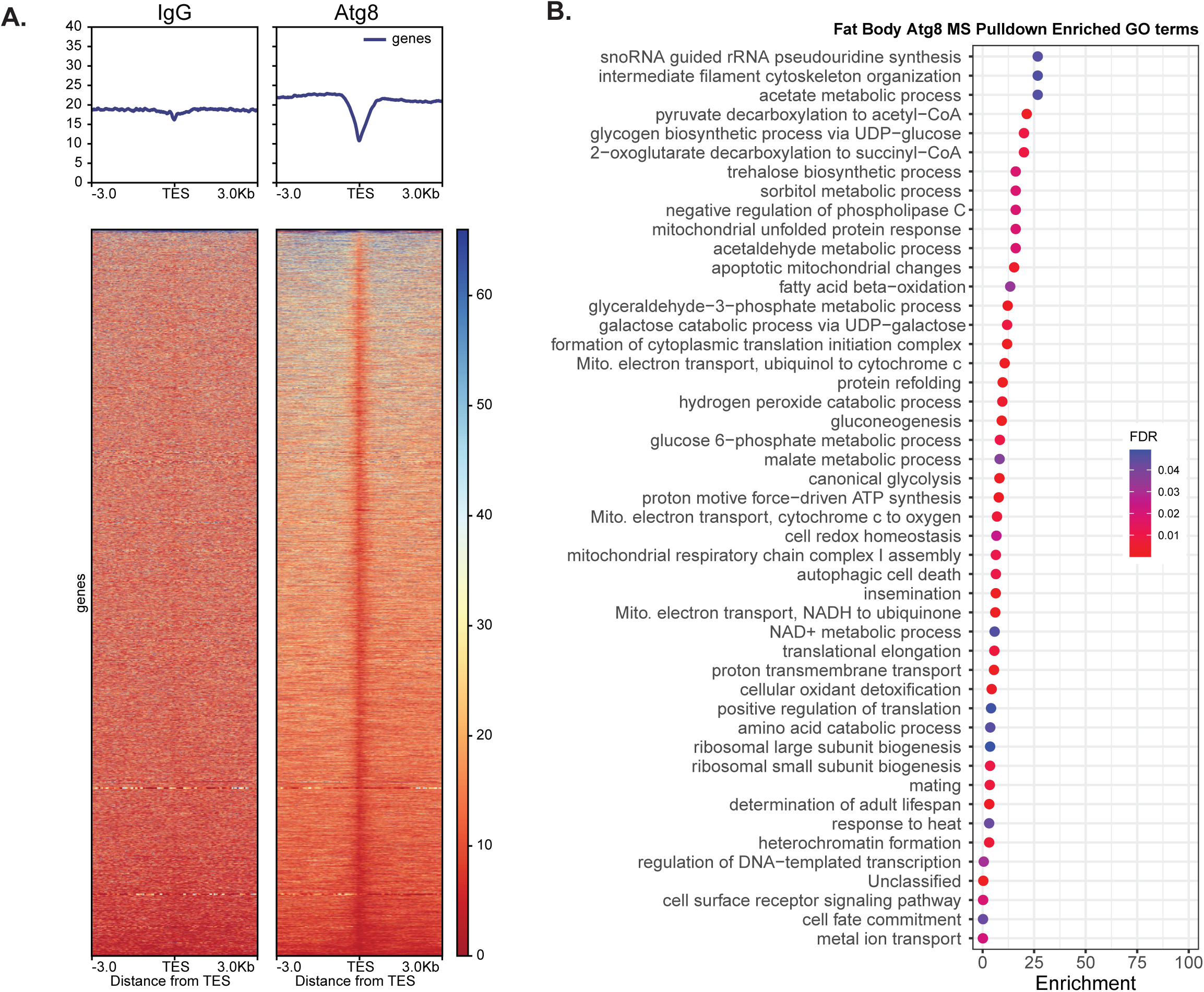
Atg8 TES occupancy and protein association. A) overview of Atg8 chromatin binding in *W1118* 7-day-old adult flies on ND aligned to 5’ transcription end sites (TES), compared to IgG negative control. The heatmap depicts the number of sequences relative to other TES in arbitrary units, with each row being a gene. Top graphs plot the mean occupancy across all TES within +/- 3kb. B) GO enrichment analysis of pathways of proteins pulled down with Atg8 from *Drosophila* fat bodies.

**Figure 3 Supplement.**
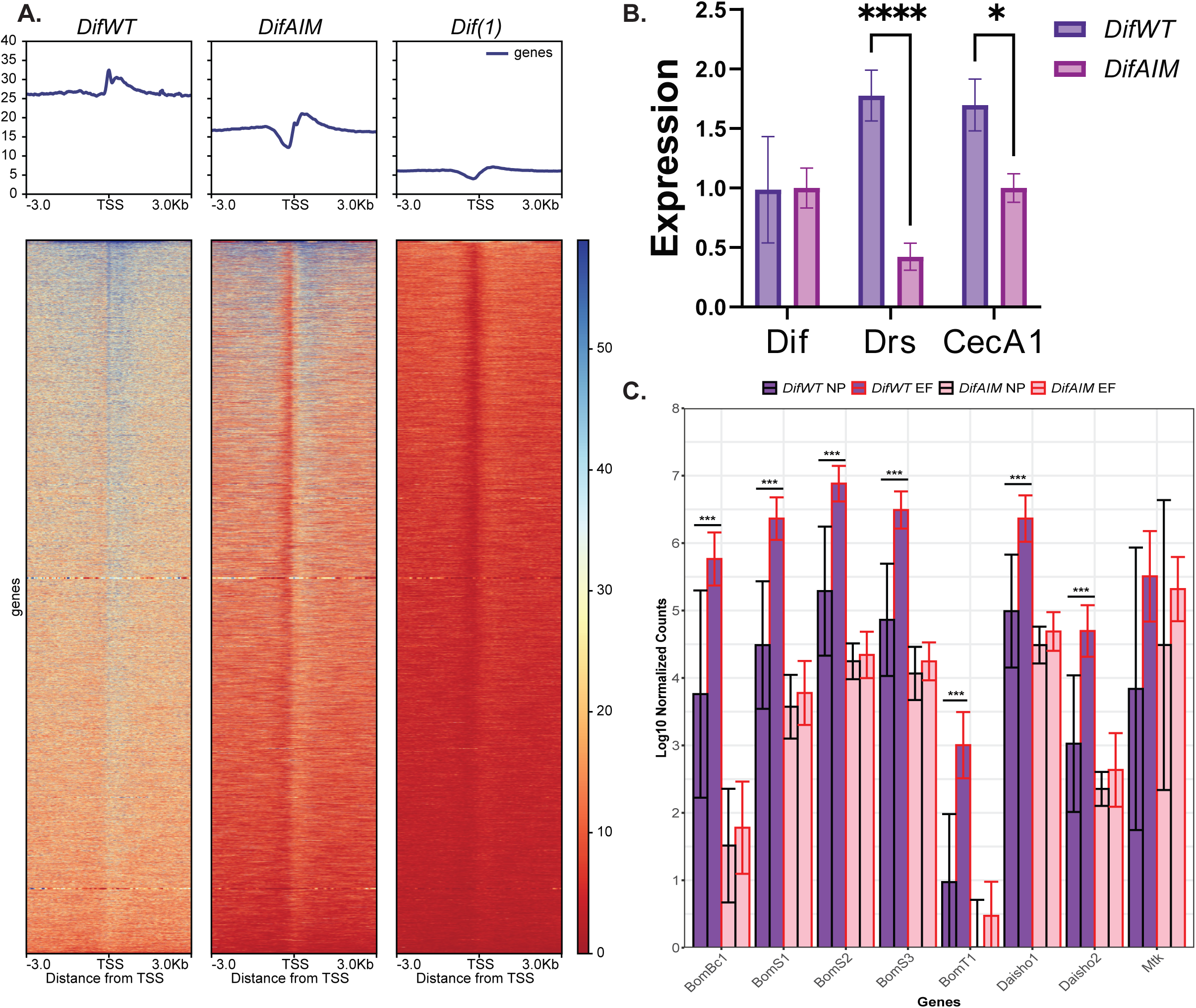
A) Full heatmap of Dif Occupancy at TSS in *DifWT*, *DifAIM*, and *Dif(1)* flies. B) qPCR analysis showing reduced basal expression of Dif target AMPs (*Drosomycin* and *Cecropin A1*) in *DifAIM* (pink) compared to *DifWT* (purple) flies. C) full list of Gram-positive AMPs enriched in RNAseq Differential Expression data in *DifWT* infected vs uninfected which are unresponsive in *DifAIM* infected vs uninfected (padj<0.05). Error bars indicate standard deviation. Significance was determined by two-way ANOVA with Bonferroni post hoc correction. *p<0.5,**p<0.1,***p<0.01,****p<0.001.

**Figure 4 Supplement.**
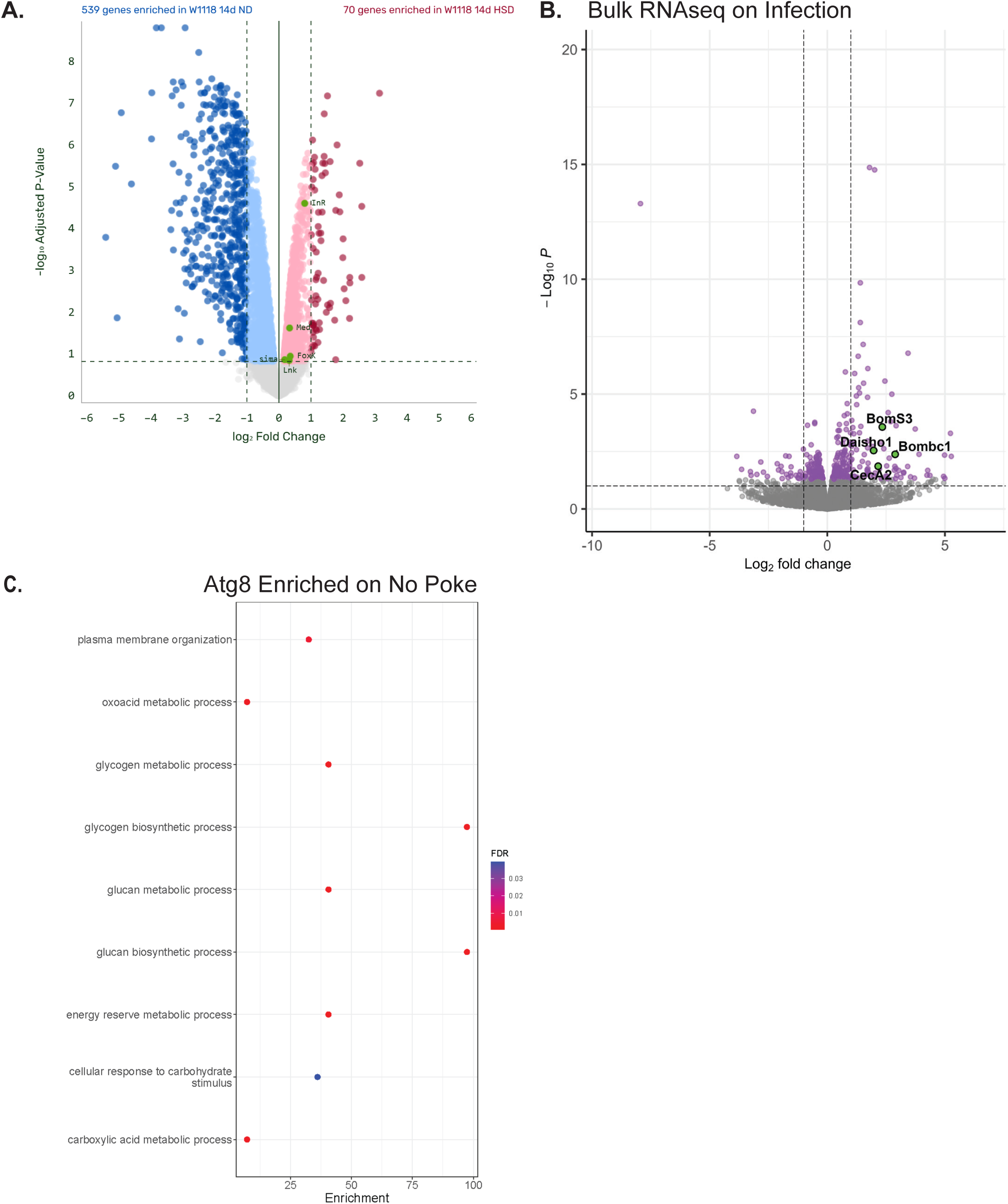
A) Volcano plot of RNAseq data of *W1118* flies in 14d ND vs HSD and B) *DifWT* flies no poke vs *E. Faecalis* 6 hours post infection. C) GO analysis of genes found to be significantly upregulated and bound by Atg8 in No Poke flies compared to *E. faecalis-*infected flies.

## Notes

### Competing Interest Statement

The authors have declared no competing interest.

### Summary of Updates

Author Shannon N. Marschall was missed in previous submission.

